# Ultra-fast super-resolution imaging of biomolecular mobility in tissues

**DOI:** 10.1101/179747

**Authors:** Helen Miller, Jason Cosgrove, Adam J. M. Wollman, Peter O’ J. Toole, Mark C. Coles, Mark C. Leake

## Abstract

Super-resolution techniques have addressed many biological questions, yet molecular quantification at rapid timescales in live tissues remains challenging. We developed a light microscopy system capable of sub-millisecond sampling to characterize molecular diffusion in heterogeneous aqueous environments comparable to interstitial regions between cells in tissues. We demonstrate our technique with super-resolution tracking of fluorescently labelled chemokine molecules in a collagen matrix and *ex vivo* lymph node tissue sections, outperforming competing methods.

---

The emergence of super-resolution imaging has had an enormous impact on biology, enabling spatial localisation of single fluorescent probes more than an order of magnitude better than the optical resolution limit of ∼250 nm, facilitating direct visualisation of dynamic molecular processes within complex biological systems^1^. Barriers to using super-resolution for quantifying rapid molecular diffusion in inter- and intra-cellular regions in tissues include poor time resolution, due to constraints imposed from limited photon emission and challenges in data interpretation due to mobility heterogeneity. Whilst elastic and interferometric scattering can overcome poor fluorophore photo-physics to enable rapid sampling they either utilise relatively large probes that exhibit steric hindrance or achieve poor specificity in heterogeneous sample environments unless used in conjunction with fluorescent labelling. Scanning fluorescence methods such as stimulated emission depletion microscopy (STED) are limited to ∼1 Hz typical frame rates with faster imaging up to ∼1,000 Hz possible by trading image quality^2^, while widefield approaches such as fast variants of photoactivatable localization microscopy (PALM)^3^ and stochastic optical reconstruction microscopy (STORM)^4^ have integration times of ∼tens of milliseconds for individual image frames with full reconstructions commonly taking several seconds. Structured illumination at best achieves several hundred Hz frame rates^5^, while high intensity illumination methods have enabled super-resolution imaging at millisecond timescales^6,7^. Sub-millisecond molecular tracking was reported previously using relatively large fluorescent bead^8^ and quantum dot probes^9^ *in vitro*, and dye labelled cholesterols in live cell membranes^10^. However, these methods encounter significant challenges in data interpretation when samples and mobility are heterogeneous, as encountered in tissues. Our method is the first, to our knowledge, to enable sub-millisecond molecular tracking using a minimally perturbative nanoscale organic dye reporter in a heterogeneous aqueous environment typical of interstitial regions between cells in tissues.

To overcome previous limitations we adapted a standard inverted epifluorescence microscope, making important modifications to facilitate sub-millisecond super-resolution tracking of rapidly diffusing fluorescently labelled biomolecules and developing bespoke software for precise quantification of underlying molecular mobility. Specifically, the optical path of the microscope was modified (Fig. 1a) to implement a broadband laser whose output was selectable over wavelengths ∼400-2,000 nm (Supplementary Fig. 1), spanning the excitation spectra of visible light and near infrared fluorophores; the beam was de-expanded using a series of lenses to generate a narrow illumination field of ∼12 μm full width at half maximum. High contrast epifluorescence images magnified to 120 nm/pixel were captured by an ultrasensitive back-illuminated EMCCD detector (860 iXon+, Andor Technology Ltd.) which could be sub-arrayed to 29x128 pixels to enable rapid frame rates of 1,515 Hz. Images were analysed using bespoke software^11^ written in MATLAB (Mathworks), enabling automated 2D sub-millisecond tracking of single fluorophores and determination of the microscopic diffusion coefficient *D* from the measured mean square displacement^12^.

**Figure 1:**
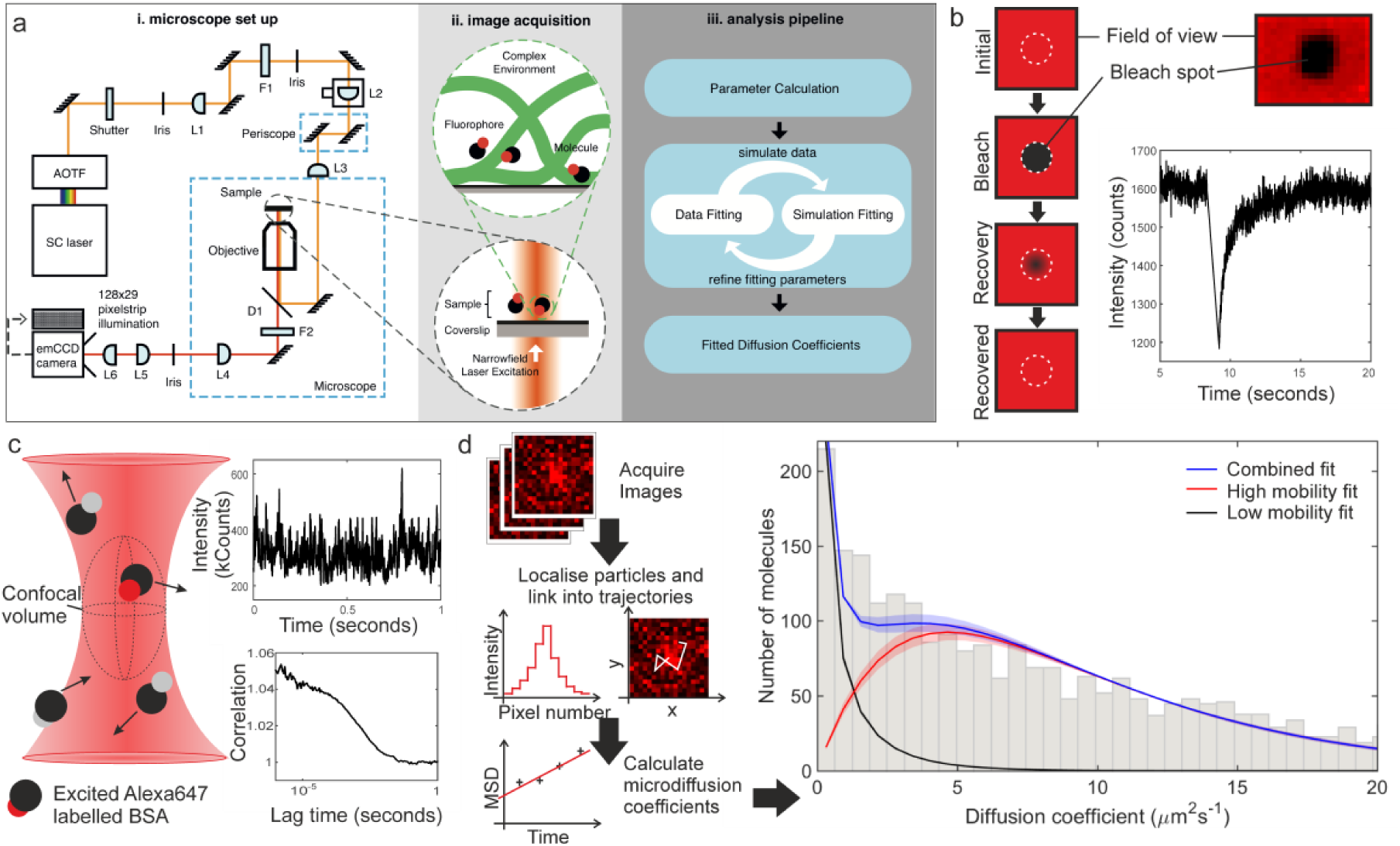
The high-speed narrowfield microscopy framework and comparison to FRAP and FCS for the test molecule, BSA-AF647 in 10% Ficoll 400. (a) High-speed narrowfield microscopy framework. (i) The bespoke fluorescence microscope. (ii) Schematic diagram of image acquisition. (iii) The analysis pipeline. (b) FRAP: schematic of technique and fluorescence intensity recovery trace. (c) FCS: Schematic of the confocal volume, section of intensity fluctuation trace and example correlation curve. (d) Single-molecule tracking: Simplified schematic of the stages in tracking and the resulting fit with error bounds of one standard deviation.

As proof-of-concept we use this scheme to quantify the diffusion of chemotactic cytokines (chemokines) CXCL13 and CCL19, key regulators of lymphocyte migration^13^. Chemokines are small proteins (∼10kDa) that bind G-protein Coupled Receptors (GPCRs) leading to polarization of the actomyosin cytoskeleton and directed migration along localised concentration gradients^14^. Due to an intricate regulatory network across broad spatiotemporal scales, direct visualisation of these molecules *in situ* is challenging. Chemokines are secreted within a dense, heterogeneous microenvironment undergoing transient interactions with cognate GPCRs and components of the extracellular matrix (ECM) before undergoing receptor-mediated scavenging or enzymatic degradation. Additionally, chemokines are heterogeneous in binding affinities and are subject to multimerisation effects, characteristics that may alter their mobility^15,16^. Hydrodynamic predications employing estimations for the Stokes radius of chemokines and the fluid environment viscosity suggest that chemokine diffusion in hypothetically homogeneous intracellular media in the absence of binding effects is rapid at ∼150 μm^2^/s, implying ∼50 s for a single molecule to diffuse across a 200 μm diameter region of lymphoid tissue. However, this estimate is likely to be a poor predictor of diffusivity as it does not account for dynamic molecular interactions encountered in dense tissues.

We investigated Alexa Fluor 647 (AF647) tagged CXCL13 and CCL19, exploring a range of sample exposure times (0.44-1.98 ms per frame), acquiring most data using 0.59 ms per frame (0.65 ms cycle time) as a compromise between sampling speed and localisation precision. We used a range of environments of increasing complexity comprising chemokines in (i) buffer alone and in the presence of the highly-branched polysaccharide Ficoll to vary the fluid environment viscosity, (ii) the presence of either surface-immobilized heparan sulphate, or a collagen gel matrix, or (iii) in an *ex vivo* native mouse lymph node environment. In all cases we were able to resolve distinct diffusing fluorescent foci of measured 2D half width at half maximum of ∼250-300 nm, consistent with single point spread function (PSF) images.

Automated foci tracking was utilised for the determination of molecular diffusivity. Foci could be tracked continuously in 2D with ∼40 nm precision In PBS buffer alone the chemokine diffusion was in general too fast to track over consecutive image frames (Supplementary Video 1), consistent with theoretical estimates for *D* >100 μm^2^/s, however applying 10% Ficoll increased the fluid viscosity by a factor of 5.6 to 0.005 Pa.s enabling single particle tracking, tested first on a non-chemokine control of AF647-tagged BSA (BSA-AF647). The results of single particle tracking of BSA-AF647 were consistent with a proportion of immobile tracks associated with the coverslip surface and a freely diffusing mobile population with *D*_*mobil*e_ = 9.3 ± 0.4 μm^2^s^-1^ (Fig. 1d and Supplementary Video 2), in agreement with theoretical expectations based on hydrodynamic modelling of BSA as a Stokes sphere of radius 3.48 nm incorporating Faxen’s law (Supplementary Note)^17^. Using the traditional molecular mobility tools of fluorescence recovery after photobleaching (FRAP) and fluorescence correlation spectroscopy (FCS) the BSA-AF647 control in Ficoll indicated *D* = 7.1 ± 0.3 μm^2^s^-1^ and 18.8 ± 0.3 μm^2^s^-1^ respectively (Fig. 1b,c). The result (Supplementary Table 1) for FRAP is significantly lower than the theoretical value, even considering Faxen’s law, temperature fluctuation and non-monomer content in BSA (Supplementary Table 2) whilst that for FCS is higher. The FRAP and FCS results differ by a factor of 2.6, in general agreement with previous results^18–20^ and often attributed to the different spatial scales of the two measurements or the high number of assumptions required in fitting FRAP data^21^. A key advantage of our rapid single particle tracking method is its ability to determine the underlying heterogeneity in the mobility of the molecular population, exemplified here by chemokines which diffuse in different environments.

Analysis of the tracking data from CXCL13-AF647 and CCL19-AF647 using step-wise dye photobleaching showed predominantly monomeric populations for each, with higher stoichiometry foci accounted for by random overlap of single dye tracks (Supplementary Fig. 1). The broad probability distributions for *D* found for both chemokines (Supplementary Fig. 2) are indicative of underlying heterogeneity in molecular mobility. As expected, FRAP and FCS failed to report on the population of bound immobilized chemokines, however, modelling the sub-millisecond tracking data as a mixture of immobilized and mobile tracks generated excellent agreement (Supplementary Table 3), corroborated through simulations of diffusing and immobilized foci using realistic signal and background noise values (Supplementary Note, Supplementary Fig. 3 and Supplementary Table 4). Our findings indicated a higher diffusion coefficient for CCL19-AF647 than for CXCL13-AF647 in the controlled environment of collagen (Fig. 2a, Supplementary Videos 3,4, and Supplementary Note), which were measured as 8.4 ± 0.2 μm^2^s^-1^ and 6.2 ± 0.3 μm^2^s^-1^ respectively. This heterogeneity is consistent with molecular mass expectations and may contribute to the formation of distinct spatial patterning profiles *in situ.*

**Figure 2:**
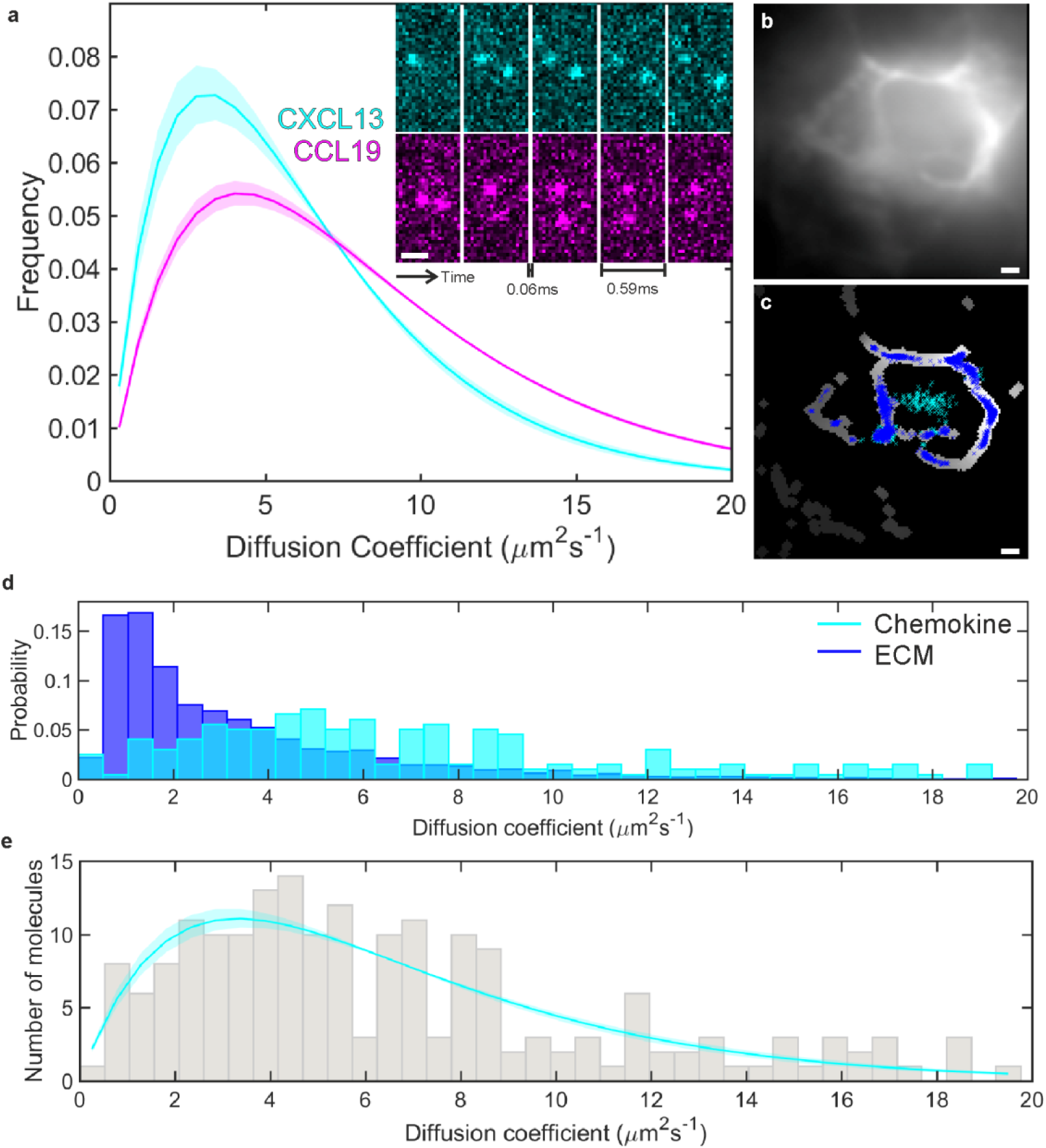
Single molecule analysis of chemokine diffusion. (a) Fitted high mobility diffusion coefficient distribution of CXCL13-AF647 (cyan) and CCL19-AF647 (magenta) in a collagen matrix (shaded areas indicate one standard deviation) with (inset) representative sub-millisecond images from these data sets. (b) Intensity average image of image acquisition to show autofluorescent ECM. (c) Areas of (b) identified as ECM by segmentation with overlaid track localisations coloured by location on ECM (blue) or elsewhere (cyan). (d) Comparison of diffusion coefficients for the ECM (blue) and chemokine (cyan) populations when tracking CXCL13-AF647 in lymph node tissue shown. (e) Distribution and fit of chemokine diffusion coefficients of CXC13-AF647 in tissue sections, shaded area indicates one standard deviation. All scale bars 1μm.

We further imaged and tracked CXCL13-AF647 in B cell follicles of *ex vivo* murine lymph node tissue sections with super-resolution precision at ∼2 ms timescales (Fig. 2b and Supplementary Videos 5,6), determining location in the tissue using FITCB220 (B cell specific marker). Auto-fluorescent extracellular matrix (ECM) components (Supplementary Note and Supplementary Fig. 3) were localised by the tracking software and were segmented to allow discrimination of tracks from the immobile ECM and the diffusing chemokine (Fig. 2b,c,d). The diffusion coefficient of mobile tracked particles was fitted with a single distribution indicating *D* = 6.6 ± 0.4 μm^2^s^-1^ (Fig. 2e).

Our novel high-speed microscopy and analysis outperforms traditional molecular mobility tools of fluorescence recovery after photobleaching (FRAP) and fluorescence correlation spectroscopy (FCS) in being able to capture diffusional heterogeneity relevant to real, complex biological systems exemplified by underlying mobile and immobile states. The high time resolution achieved with our system enables rapid diffusion to be quantified in heterogeneous aqueous environments typical of interstitial regions between cells in tissues, whilst still retaining super-resolution spatial precision and single-molecule detection sensitivity, enabling new insight into complex systems. Our system is compatible with traditional widefield light microscopes as opposed to requiring expensive and dedicated super-resolution setups; this accessibility bodes well for establishing a significant future impact investigating multiple biological systems.

## Methods

Methods and associated references are available in the online version of the paper.

*Note: Any Supplementary Information and Source Data files are available in the online version of the paper.*

## Acknowledgments

Supported by the Biological Physical Sciences Institute, MRC grants MR/K01580X/1 (P.O.T. and M.C.L), MC_PC_15073 (M.C.C. and M.C.L) and BBSRC grant BB/N006453/1 (A.J.M.W. and M.C.L.). J.C. is supported by a studentship from the Wellcome Trust 4-year PhD programme (WT095024MA): Combating Infectious Disease: Computational Approaches in Translation Science. The authors thank Jo Marrison and Andrew Leech (Bioscience Technology Facility, University of York) for technical assistance with FCS and FRAP microscopy, and for SEC-MALLs respectively, Chris Power (Carl Zeiss Microscopy) for help with FCS, and Anne Theury for providing lymph node tissue sections.

### AUTHOR CONTRIBUTIONS

H.M. built the bespoke fluorescence microscope, overseen by M.C.L.; J.C. prepared biological samples overseen by M.C.C. H.M. and J.C. performed the imaging; H.M. analysed the data from all experiments with input from A.W., M.C.L. H.M. ran simulations of fluorescence data on code adapted from A.W. P.O.T. oversaw FCS and FRAP microscopy. H.M., A.W. prepared the figures with input from all authors. H.M., J.C, and M.C.L. wrote the manuscript with input from all authors.

### COMPETING FINANCIAL INTERESTS

The authors declare no competing financial interests.

## METHODS

### 1. Reagents

The human chemokines (CCL19 and CXCL13) singly labelled with the far-red fluorescent tag AF647 (CAF-06 and CAF-12, Almac) were stored in water at 222 μg/ml. Collagen samples contained type I collagen extracted from rat tails^22^ diluted in PBS to 3.3 mg/ml and chemokine at 111 ng/ml; samples were set to pH7 with the addition of NaOH. BSA labelled with 3-6 AF647 was purchased from Thermo Fisher Scientific Inc. Ficoll 400 (Sigma-Aldrich) was diluted in PBS at 0.1g/ml to create a 10% solution of viscosity 0.005 Pa.s at room termperature^23^.

### 2. Preparation of collagen matrix in tunnel slides

Samples for fluorescence microscopy were prepared in tunnel slides formed by placing two parallel lines of double-sided tape on a standard microscopy slide around 5 mm apart. A plasma cleaned coverslip was placed on top and carefully tapped down (avoiding the imaging area) to create a water-tight tunnel.

For imaging in a collagen matrix tunnel slides were cooled to 4°C before addition of collagen and fluorescently labelled chemokines, and then incubated at 15°C for 30 min, followed by an additional 30 min incubation at 37°C. The collagen matrix was visualised using second harmonic imaging^24,25^.

Immobilised chemokine samples were prepared by incubating a plasma-cleaned coverslip in heparan sulphate^26^ (50 mg/ml) (Sigma-Aldrich) in PBS for 30 min. Coverslips were washed with PBS and air dried for 30 min before tunnel slide assembly. 10 nM fluorescently labelled chemokine solution in PBS was introduced to the tunnel slide and incubated in a humidity chamber for 15 min at 20°C. Excess unbound chemokine was removed with a PBS wash.

### 3. SHIM imaging

Second harmonic imaging microscopy (SHIM) was performed on a Zeiss LSM 780 MP with a Zeiss invert microscope. Excitation at 900 nm wavelength (Coherent Ultra Laser) through a plan-apochromat 63x/1.4 oil objective lens was incident on the sample. Up converted light was collected via a 485 nm short pass filter onto a non-descanned detector.

### 4. Preparation of Lymph Node Tissue Sections

6-8 week old wild type mice (C57BL/6) were housed in BSF at the University of York. All experiments conformed to the ethical principles and guidelines approved by the University of York Institutional and Animal Care Use Committee. Popliteal Lymph Nodes were removed and excess fat or connective tissue removed with forceps. Samples were transferred to optimal cutting temperature medium (OCT, Tissue-Tek, Sakura Finetek) and snap frozen on dry ice samples and sectioned using a cryostat. 10 μm thick sections were cut and collected onto poly-L-Lysine coated microscope slides. Sections were dried overnight in the dark then stored at −20°C.

Before use, lymph node sections on poly-L-lysine slides were incubated at room temperature for 30 min. Sections were hydrated in PBS for 5 min then air dried. Wax ImmEdge pen (Vector Laboratories) was used to draw a hydrophobic circle around each section to retain liquid on the section during staining. Sections were incubated in a blocking buffer of PBS 5% goat serum (Sigma) at room temperature for 1 hour. To determine where B-cells were located in the tissue we used the marker B220, a protein expressed on the surface of all murine B cells. After blocking, sections were incubated with an FITC conjugated antibody (RA3-6B2, purchased from eBioscience) that binds specifically to B220 diluted 1:200 in blocking buffer for 1 hour at room temperature. Samples were washed with PBS for 3 × 5 min and 1μM of CXCL13-AF647 added to the slides. Slides were left to incubate overnight at 4°C after which slides were washed for 30 s in PBS and a coverslip (thickness 0.13-0.17mm, Menzel Gläser) mounted on top. Slides were then sealed and imaged.

### 5. Stokes model of diffusion

For a small sphere the diffusion coefficient is given by

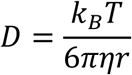

Where k_B_ is the Boltzmann constant, T is room temperature, η is the dynamic viscosity of the media and *r* is the radius of a sphere. The theoretical value of the diffusion coefficient of BSA in 10% Ficoll 400 was calculated using the Stokes radius of BSA of 3.48nm ^27^.

### 6. FCS and FRAP microscopy

FCS and FRAP experiments were performed on a Zeiss LSM 880 microscope, using a GaAsP detector. Samples were prepared in MatTek glass bottom petri dishes (1.5 coverglass, MatTek corporation) and illuminated with a 633 nm wavelength laser.

For FCS the confocal volume was measured using a calibration sample of BSA-AF647 diffusion in water and constraining the diffusion coefficient to be 59 μm^2^s^-1^ ^28^; this allowed the structural parameter, *s*, to be fixed at 6.6. Three repeats of ten experiments were conducted, traces indicating the presence of multimeric clumps or proximity to the surface were excluded. Autocorrelation traces, G(τ), to account for transient dark states and translational diffusion were fitted with the equation:

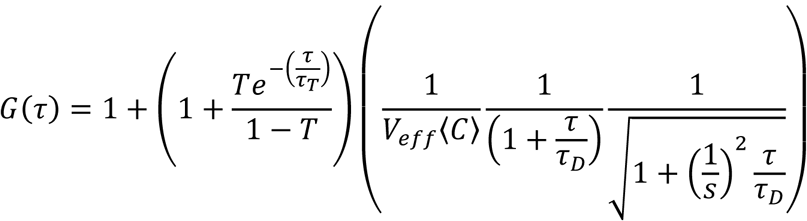

Where T is the triplet fraction, τ_T_ is the time constant of the dark state, *τ*_*D*_ is the time constant of translation across the confocal volume, *V*_*eff*_, and <*C*> is the average concentration. The diffusion coefficient, *D*, was calculated from *τ*_*D*_ via the equation:

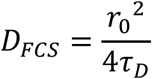

Where *r*_*0*_ is the spot width (0.322 μm). For FRAP microscopy a square region of half-width 4.9 ± 0.1μm, *ω*, (measured with an immobilised sample) was bleached with the 633 nm wavelength focused laser. To measure the diffusion coefficient of BSA-AF647 thirty recovery traces (intensity (*I*) vs time (*t*)) were acquired and fitted in the Zeiss Zen software with the single exponential equation:

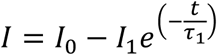

Where the initial intensity, *I*_*0*_, drop in intensity, *I*_*1*_, and the decay constant *τ*_1_, from which the diffusion coefficient, *D*, is calculated via:

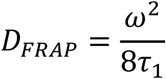

### 7. Fluorescence microscopy

Bespoke fluorescence microscopy was performed on an inverted microscope body (Nikon Eclipse Ti-S) with a 100x NA 1.49 Nikon oil immersion lens and illumination from a supercontinuum laser (Fianium SC-400-6, Fianium Ltd.), controlled with an acousto-optical tunable filter (AOTF) to produce excitation light centred on wavelength 619 nm (Supplementary Fig. 1). A 633 nm dichroic mirror and 647 nm long-pass emission filter were used beneath the objective lens turret to exclude illumination light from the fluorescence images. The sample was illuminated with narrowfield excitation of 12 μm FWHM and an intensity of 2,300 W.cm^-2^. Images were recorded on an EMCCD camera (860 iXon+, Andor Technology Ltd) cooled to −80°C. 128×128 pixel images were acquired with 1.98 ms exposure times and 128 × 29 pixel image strips were acquired with 0.59 ms exposure times, both for 1,000 frames at the full EM and pre-amplifier gains of 300 and 4.6 respectively.

### 8. Particle Tracking and calculation of diffusion coefficients

All image data were recorded into .tiff files and analysed in bespoke Matlab software. Single fluorescent molecules were identified and processed into super-resolution tracks using ADEMS code^11^. The microscopic diffusion coefficient was calculated for each tracked particle from the gradient of a linear relation fitted to the first four steps in a track. The microscopic diffusion coefficient distributions comprised an immobile fraction which had non-specifically adhered to the plasma-cleaned coverslip and a diffusive fraction. Microscopic diffusion coefficients were binned into histograms with bin width given by the localization precision of the immobilised (heparan sulfate) data. The probability distribution of diffusion coefficients was modelled by a gamma distribution^29–32^:

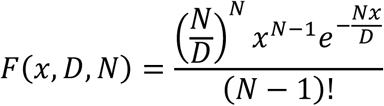

Where *N* is the number of independent steps in a track and *D* is the true diffusion coefficient. The histogram data was fitted iteratively with a two gamma distribution to account for the mobile and immobile fractions. Initial fitting constraints conserved the number of tracks and assumed the number of independent steps in a track was 4 or less, giving a first estimation of the diffusion coefficients. Then fluorescence microscopy data with the found diffusion coefficients was simulated with and without noise, tracked, and the distribution of diffusion coefficients was fitted with the same constraints as the actual data. Fitting parameters were refined based on the results of fitting to the simulated data, and the experimental data was fitted with the refined constraints. The process of simulating the data, fitting the simulation to refine the constraints and fitting the experimental data was repeated until the simulation represented the experimental data and the fit to the simulation data converged to the diffusion coefficient values simulated.

For immobilized spots the *N* value was less than 1, implying that the steps are not independent. This is expected for immobile molecules as the localisation precision uncertainty is larger than the diffusion distance. For mobile spots *N* was fixed at two as there are two steps which do not contain any common localisations when only the first four steps of a track are used.

### 9. Simulation of fluorescence microscopy data.

Image datasets were simulated in bespoke Matlab software at given diffusion coefficients using foci intensity, background intensity and foci density data from real images. Foci are created at random locations in the image frame with intensity randomly chosen from a localization in an experimental dataset. The distance travelled by a foci is taken from a Gaussian distribution centred on the spot location with the width of the mean square displacement of a particle with the simulated diffusion coefficient. To incorporate photobleaching and other effects causing truncation of trajectories foci were randomly reassigned to a new location 10% of the time. The resulting image stack was used for no-noise simulations. Readout noise in the detector was incorporated in the simulations by the addition of zero mean Gaussian white noise, the intensity of which depended on the local intensity.

### 10. Bootstrapping

Errors on the found values of the diffusion coefficients from FRAP, FCS and single molecule tracking were found by bootstrapping^33,34^. In this method, a randomly chosen 80% of the data is fitted in the same way as the entire data set and the standard deviation on each parameter from ten repeats of this process is taken as the error on each fit parameter found from 100% of the data.

### 11. Imaging in tissue sections

To determine where B-cells were located in the tissue we used the marker B220, a protein expressed on the surface of all murine B cells. Tissue sections were stained with a FITC-conjugated antibody that binds B220. Tissue sections were subsequently imaged at low (1.2 μm/pixel) magnification with green illumination (wavelength 470 nm, 12 μm FWHM, intensity of 875 W.cm^-2^) to determine the location of the B cell follicles, before switching to high (120 nm/pixel) magnification and red illumination to image chemokines in these areas.

### 12. Segmentation of tissue sections images

Image acquisitions in tissue contain regions of autofluorescent ECM (see Supplementary Note and Supplementary Fig. 3) which are identified by the tracking software. These images must be segmented to identify tracks due to fluorescently labelled chemokine or ECM. Intensity averages of the image acquisition were top hat filtered with a structuring element of radius 4 pixels. The resulting image was converted to binary form using a threshold defined by Otsu’s method and the binary image used to enhance the extracellular matrix regions of the original image. Small holes in the thresholded region were filled by sequential erosion and dilation with a disk of radius 2 pixels as the structuring element.

### 13. Software availability

All our bespoke software developed is freely and openly accessible via https://sourceforge.net/projects/york-biophysics/

## Supplementary notes, figures and legends

### Single-molecule Imaging

Single-molecule imaging was performed using a supercontinuum laser source in combination with an AOTF for wavelength selection. This produces a distinct spectral profile. The illumination profiles used for FITC and AF647 are shown in Supplementary Fig. 1a,b.

The presence of single molecules was determined by the observation of stepwise photobleaching steps. Four examples are shown in Supplementary Fig. S1c.

Photoblinking behavior of a single molecule shows defined steps in the intensity versus time trace between ‘on’ and ‘off’ states. An example of this is shown in Supplementary Fig. S1d, where the spot location is tracked through time. When the molecule is ‘on’ the localisations show good clustering, although they are more widely dispersed during the off periods. The intensity versus time trace is shown with sample images from the on and off times. From the manufacturers specifications BSA-AF647 was expected to be labelled with between 3-6 AF647 dye molecules and the chemokines were expected to be singly labelled. Only single molecule bleaching steps were observed in the chemokine data.

**Supplementary Figure S1:**
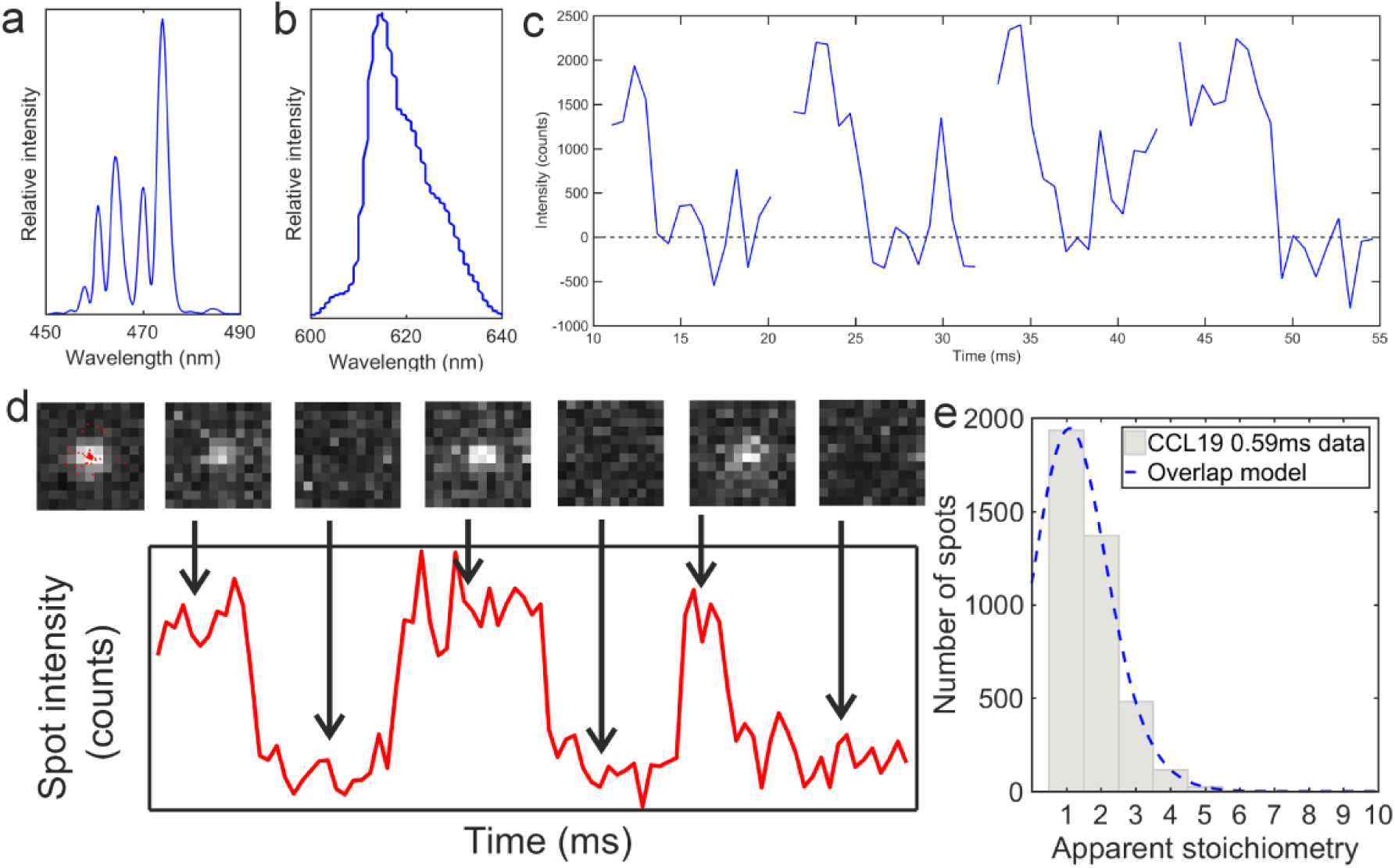
Single molecule imaging. Wavelength spectra of illumination light used for excitation of (a) FITC and (b) AF647. (c) 4 sample stepwise bleach traces from AF647 labelled CCL19. (d) Tracking of blinking AF647: localisations and intensity over time with sample images from the acquisition. (e) Distribution of apparent CCL19 foci stoichiometry (grey) overlaid with the predicted distribution based on randomly overlapping PSFs (blue).

In populations of single molecules higher intensities are often seen. These can either be real higher stoichiometry multimers, or caused by random overlap. The maximum number of detected foci in one frame (15 foci) was used in a previously developed model which assumes a Poisson distribution for nearest neighbour foci distances^35^.

This analysis indicates an 18% probability for random single foci overlap. The predicted apparent stoichiometry of foci was obtained by convolving the intensity distribution of a single molecule of AF647 with the Poisson distribution resulting from the probability of random overlap. The overlap model is statistically identical to the observed distribution below apparent stoichiometry of 6 AF647 molecules per foci (p<0.05, Pearson’s χ^2^ test). A small proportion of less than 5% of foci have a higher apparent stoichiometry than that predicted from the random overlap model which may be due to autofluorescence not accounted for the in overlap model.

### Diffusion coefficients of BSA-AF647

The diffusion coefficients of BSA-AF647 in 10% Ficoll 400 were used as a calibration for FCS, FRAP and single molecule imaging. The values found for the diffusion coefficients by these methods are summarised in Supplementary Table 1 with the number of traces used for each measurement.

**Supplementary Table S1:** Measurements of the diffusion coefficient of Alexa-647 labelled BSA in 10% Ficoll 400. Variation on the theoretical value is due to a potential ± 2°C temperature change in the laboratory.

| Condition | Diffusion coefficient ($\mu\text{m}^2\text{s}^{-1}$) | Number of measurements |
| --- | --- | --- |
| **Theoretical with Stokes radius 3.48nm** | 12.3 ± 0.1 | |
| **FCS** | 18.8 ± 0.3 | 27 traces |
| **FRAP** | 7.1 ± 0.3 | 30 repeats |
| **Single-molecule tracking** | 9.3 ± 0.4 | 2,608 tracks (fitted 1,113 mobile tracks) |

Distances probed by narrowfield fluorescence microscopy are within a few hundred nanometres of the coverslip, and whilst this distance is large compared with the radius of a single molecule of AF647 tagged chemokine or BSA there will still be a boundary effect in the solution of increased viscosity, which can be modelled by Faxen’s law^36^:

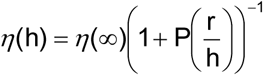

where *η*^(∞)^ = dynamic laminar-flow viscosity in free solution, and

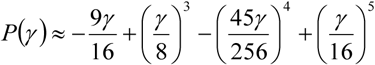

to a 5th-order approximation.

This is found to give *D* values in agreement with the single-molecule tracking measurements for particles < 10 nm from the coverslip surface.

Most molecules will be unlikely to be this close to the surface; the known multimerisation of BSA 37 should also be considered. Size Exclusion Chomatography - Multi-Angle Laser Light Scattering (SEC-MALLS) was used to examine the monomer/dimer population of the BSA-AF647 used in the comparative experiments. A ratio of approximately 15:2 monomer:dimer molecules was found (see Supplementary Table 2).

The experimental system for SEC-MALLS experiments comprised a Wyatt HELEOS-II multi-angle light scattering detector and a Wyatt rEX refractive index detector attached to a Shimadzu HPLC system (SPD-20A UV detector, LC20-AD isocratic pump system, DGU-20A3 degasser and SIL-20A autosampler). 100 μl of 2.5 mg/ml BSA-AF647 sample was run at 0.5 ml/min flow rate at room temperature through superdex S200 columns (G.E. Healthcare) for 60 min in PBS running buffer. Peaks were integrated using Astra V software and the Zimm fit method with degree 1; a value of 0.183 was used for protein refractive index increment (dn/dc).

**Supplementary Table S2:** Results of mass analysis on refractive index determined elution profile. Note, the total does not sum to exactly 100% since there is a small tail proportion not included in monomer or multimer peaks.

| Peak | Elution time from (min) | Elution times until (min) | Mass (µg) | % of total mass |
| --- | --- | --- | --- | --- |
| Monomer | 26.02 | 31.00 | 189.7 | 74.0 |
| Dimer | 22.02 | 25.98 | 48.5 | 18.9 |
| Higher oligomer | 18.00 | 21.98 | 16.5 | 6.4 |
| Total | 17.50 | 32.00 | 256.4 |  |

Ignoring the higher oligomer population which are clearly visible as large clumps of fluorescent molecules, we assume a 15:2 mixture of monomer and dimer are present in our BSA-AF647 sample. Assuming the Stokes radius of the dimer will be approximately double the Stokes radius of the monomer, 6.96 nm, the expected diffusion coefficient of the dimer is 6.2 μm^2^s^-1^. Weighting the average of the diffusion coefficients according to the relative monomer and dimer populations gives an average diffusion coefficient of 11.6 μm^2^s^-1^, closer to the value observed by single molecule tracking.

### Diffusion coefficients of chemokines in collagen

Theoretically, the diffusion coefficients of the chemokines can be calculated using the Einstein-Stokes equation, and are given in Supplementary Table 3. For these calculations the radius of the sphere was calculated assuming that the molecule is a globular protein of density 1.35 g.cm^-3^ ^38^.

To perform single-molecule distribution fitting, a histogram of the data is required with appropriate bin size. We used the localisation precision of heparan sulfate immobilised chemokines labelled with AF647 of ∼40 nm (Supplementary Video 6,7), from the low diffusion coefficient peaks in the heat maps shown in Supplementary Fig. 2a,b.

**Supplementary Fig. S2:**
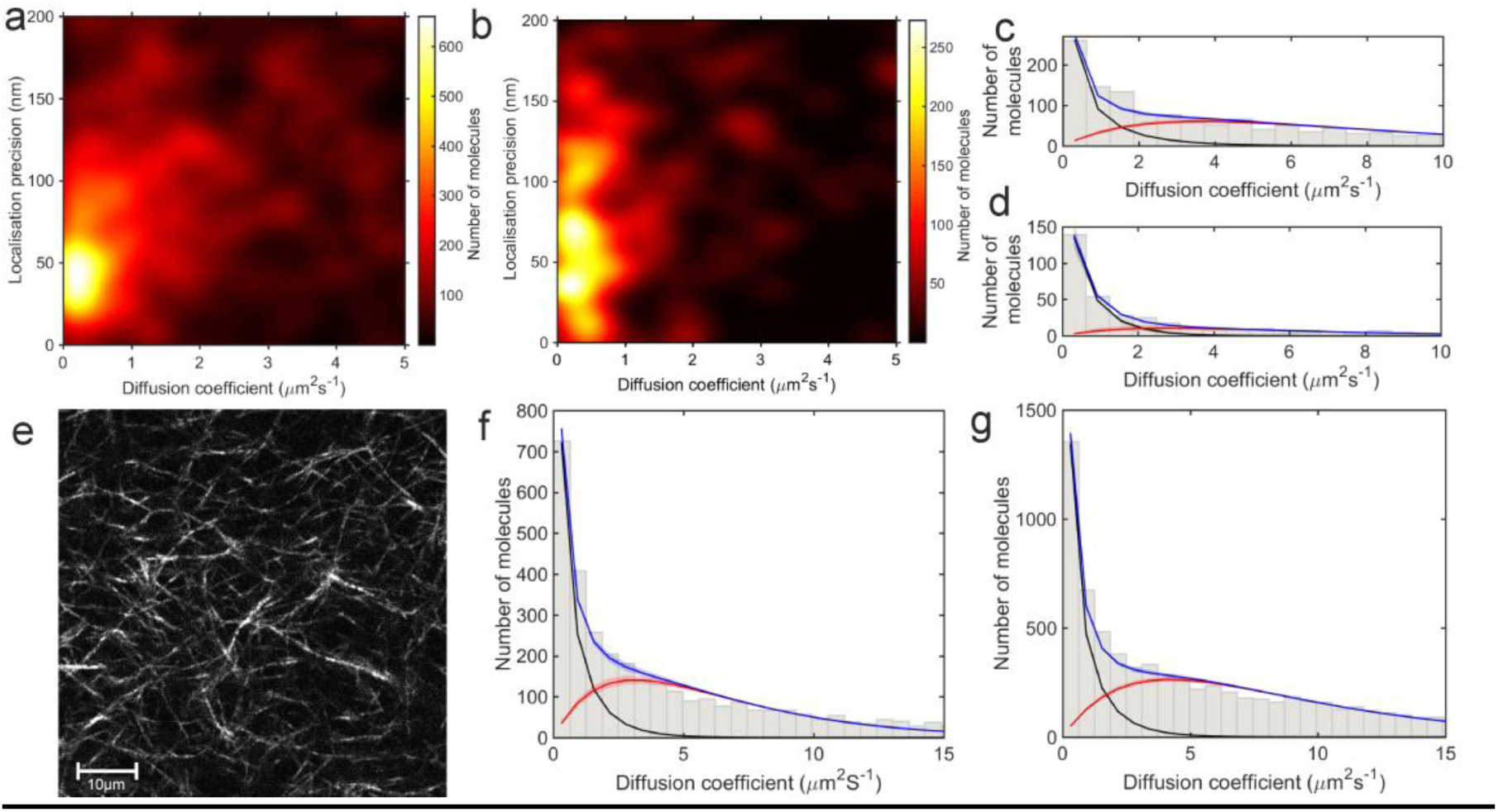
Measuring diffusion coefficients of chemokines in collagen. Heat maps of the localisation precision for heparan sulfate immobilized (a) CXCL13-AF647 and (b) CCL19-AF647, the corresponding microscopic diffusion coefficient distributions are shown in (c,d); the fitted populations are shown in black for the immobile, red for the mobile and blue for the combined fit. (e) 2D SHIM image of collagen network. Scale bar 10μm. (f,g) Microscopic diffusion coefficient distributions of CXCL13-AF647 and CCL19-AF647 in collagen; the colours are the same as (c,d).

The distributions of the microscopic diffusion coefficients of heparan sulfate immobilised chemokines are shown in Supplementary Fig, 2c,d, showing a high immobile peak and small mobile population.

The values of the diffusion coefficients were determined in collagen, the structure of which was checked with SHIM imaging (see Supplementary Fig. 2e). The fibril diameters observed are in agreement with those seen by Chen *et al.* ^25^, and show qualitatively similar structure.

The values of the diffusion coefficients of CXCL13 and CCL19 in collagen found by a two gamma distribution fit to the single molecule tracking data are given in Supplementary Table 3, and the distributions for each chemokine showing the mobile and immobile populations are shown in Supplementary Fig. 3(f,g). The relative size of the mobile and immobile populations cannot be accounted for as immobile populations were photobleached to enable imaging of highly diffuse mobile populations.

**Supplementary Table S3:** Diffusion coefficients of CXCL13 and CCL19 in collagen found by fitting a two gamma distribution to single molecule tracking data.

| **AF647**<br>**labelled**<br>**chemokine** | **Theoretical**<br>**diffusion**<br>**coefficients in**<br>**water ($\mu\text{m}^2\text{s}^{-1}$)** | **Fitted**<br>**diffusion**<br>**coefficient**<br>**($\mu\text{m}^2\text{s}^{-1}$)** | **Error**<br>**($\mu\text{m}^2\text{s}^{-1}$)** | **Number of**<br>**highly**<br>**mobile**<br>**tracks** | **$R^2$ of**<br>**combined**<br>**fit** |
| --- | --- | --- | --- | --- | --- |
| **CXCL13** | 149 | 6.2 | 0.3 | 1930 | 0.980 |
| **CCL19** | 146 | 8.4 | 0.2 | 4859 | 0.984 |

### Results of data simulation

Iterative cycles of simulation and experimental data fitting were used to determine the initial parameters for fitting and the form of the fitting functions. All simulations were run and fitted with and without the addition of Gaussian white noise. The first simulated values were chosen by fitting the experimental data with a two gamma distribution model to account for two diffusive populations.

Initially two distributions; 1.6 μm^2^s^-1^ and a 50:50 mixed population of 1.6 μm^2^s^-1^ and 10 μm^2^s^-1^ were simulated, and fitted with 1,2 and 3 gamma distribution functions with 4 independent steps, and the χ^2^ goodness of fit parameter was evaluated. χ^2^ accounts for the number of free parameters in the fit, and is used to determine if decreasing residuals are caused by increasing the number of free parameters. For the one component distribution the one gamma fit gave the lowest χ^2^, and for the two component distribution the two gamma fit gave the lowest χ^2^ value, as expected. Applying these three models to the experimental collagen and heparan sulfate immobilised chemokine data gave the lowest χ^2^ for a two component fit, except for CCL19 in heparan sulfate, which had a very low population of mobile tracks and was well fitted by a single component distribution. From this, it was determined that a two component distribution should be fitted to the experimental data.

The number of independent steps is usually the same as the number of steps in gamma distribution fitting of microscopic diffusion coefficients, but when the temporal resolution is increased, the localisation precision is decreased and steps containing the same localisations are no longer independent. To investigate the number of independent steps in the data, simulations of 1.6 μm^2^s^-1^ and 10 μm^2^s^-1^ were made separately, and fitted with a single component gamma distribution where the number of independent steps was allowed to vary. The results (Supplementary Table 4) give this value to be around two, in line with expectations of consecutive steps containing the same localisation and not being independent, reducing the number of steps by half.

**Supplementary Table S4:**
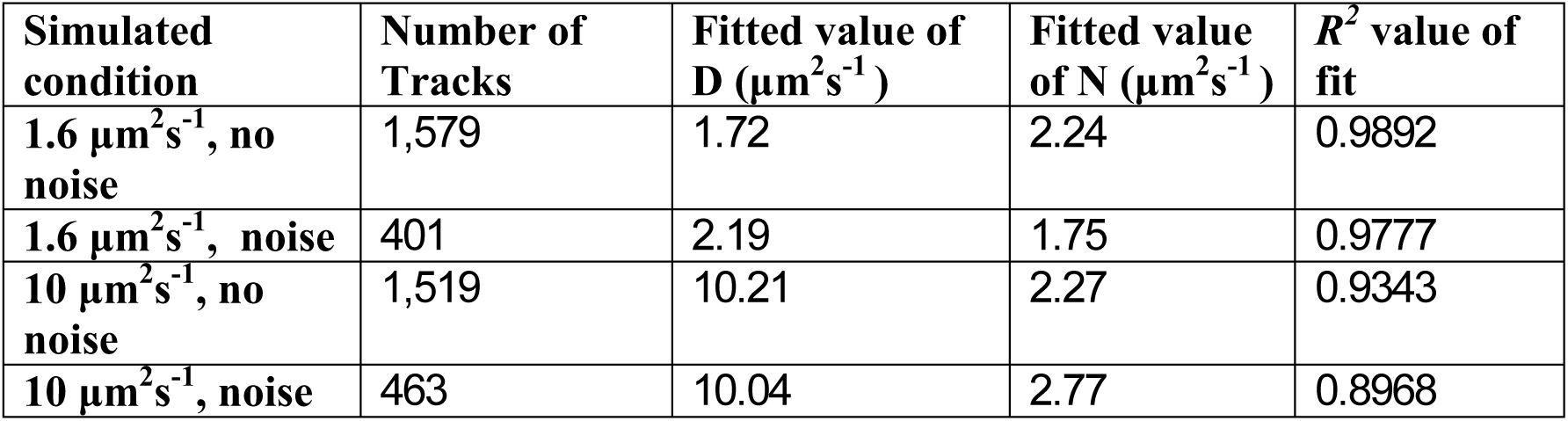
Results of one gamma distribution fitting to simulated single diffusion coefficient distributions. Noise or no noise refers to the presence of Gaussian white noise proportional to the intensity in the simulation.

| Simulated condition | Number of Tracks | Fitted value of $D$ ($\mu\text{m}^2\text{s}^{-1}$) | Fitted value of $N$ ($\mu\text{m}^2\text{s}^{-1}$) | $R^2$ value of fit |
| --- | --- | --- | --- | --- |
| $1.6 \mu\text{m}^2\text{s}^{-1}$, no noise | 1,579 | 1.72 | 2.24 | 0.9892 |
| $1.6 \mu\text{m}^2\text{s}^{-1}$, noise | 401 | 2.19 | 1.75 | 0.9777 |
| $10 \mu\text{m}^2\text{s}^{-1}$, no noise | 1,519 | 10.21 | 2.27 | 0.9343 |
| $10 \mu\text{m}^2\text{s}^{-1}$, noise | 463 | 10.04 | 2.77 | 0.8968 |

The 1.6 μm^2^s^-1^ data was simulated because this was found as an early result of fitting to the experimental data. However, the experimental data shows a peak in the first bin width, not seen in the simulation of 1.6 μm^2^s^-1^ data. Truly immobile data was simulated, and gave a peak in the first bin of the histogram when put into bins with the width of the localisation precision (see Supplementary Fig. 3).

**Supplementary Figure S3:**
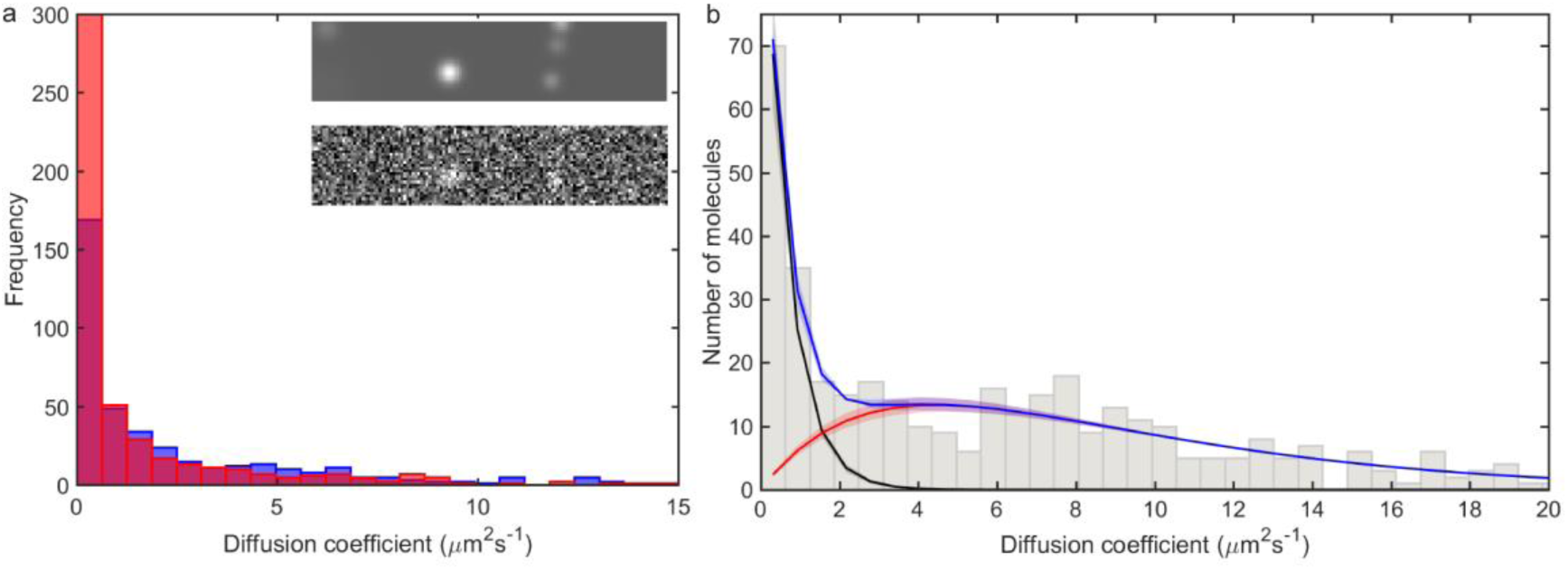
Simulations of chemokine data. (a) Distribution of simulated 0 μm^2^s^-1^ data with (blue) and without (red) Gaussian white noise. The red peak is at ∼1200, two sample images from simulations with and without Gaussian white noise are shown inset. (b) Microscopic diffusion coefficient distribution from a simulation of 0 and 9 μm^2^s^-1^ data with Gaussian white noise, Fitted populations are shown in black for the immobile, red for the mobile and blue for the combined fit.

This suggests that the low mobility population in the experimental data is immobile at the level of the localisation precision. Fitting the distribution of simulated 10 μm^2^s^-1^ data with a single gamma distribution gave a value of *N* less than one, and requires the fit applied to the experimental data to include a different value of *N* in the distribution fitted to each population; with the value of *N* being less than one for the low mobility population, and two for the higher mobility population.

Applying this fit, with the constraint that the diffusion coefficient of the immobile population must fall within the first bin of the histogram, gave the fitted experimental diffusion coefficients. To qualitatively compare the simulated and experimental data, a mixed simulation of 0 and 9 μm^2^s^-1^ data with Gaussian white noise was performed and fitted in the same way, giving a diffusion coefficient for the mobile peak of 8.9 ± 0.4 μm^2^s^-1^. The distribution is qualitatively similar in profile to the data for CCL19-AF647 in collagen (see Supplementary Fig. 3b).

### Tissue data controls

To ensure tissue measurements were performed in B cell follicles, FITC-labelled antibodies were used to detect the B cell marker in the lymph node tissue sections. Two areas, identified as B cell follicle and not B cell follicle are shown in supplementary figure S4.

Within B cell follicles, extracellular matrix components are autofluorescent in the far red, and were segmented during diffusivity analysis of tissue section data (Fig. 2b,c). When the same analysis was performed on control tissue sections prepared by the same protocol except without the addition of CXCL13-AF647 similar autofluorescent structures were seen and could be segmented (see Supplementary Fig. 3). The lack of fluorescent localisations in the gap in the extracellular matrix in the control is further evidence that the mobile population observed in the data is CXCL13-AF647.

**Supplementary Figure S4:**
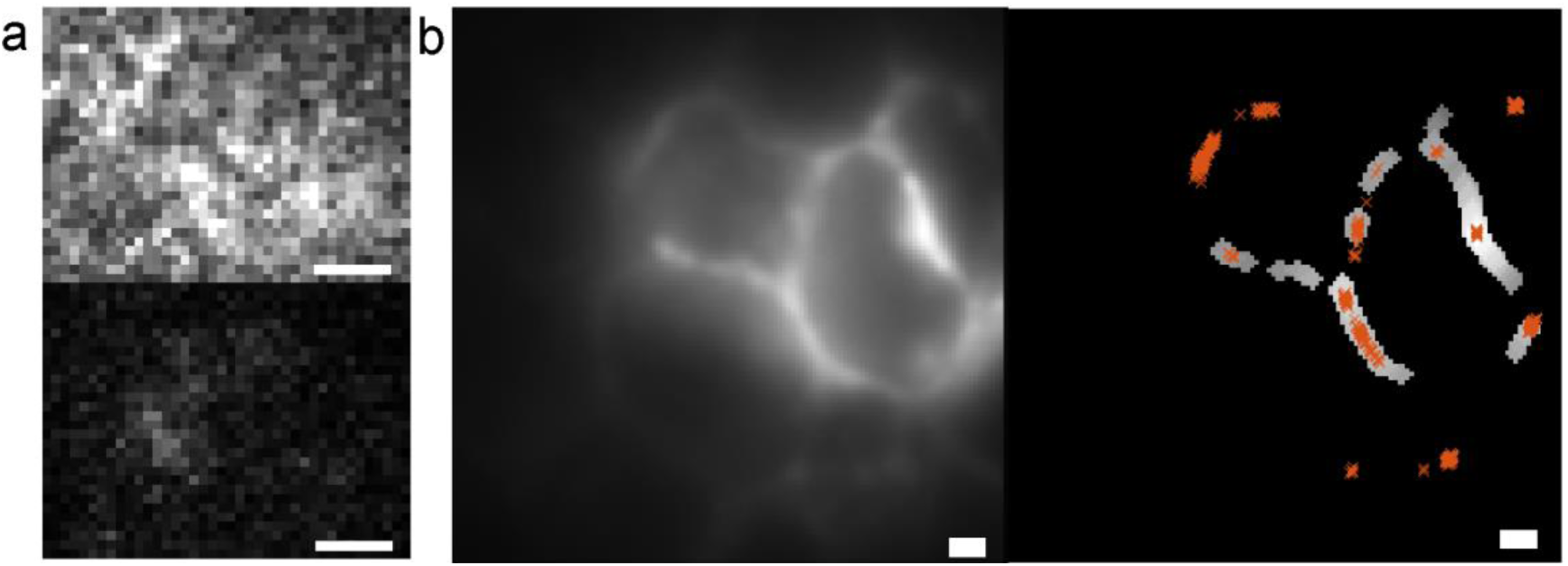
Lymph node tissue section controls. (a) Images of two regions of lymph node tissue displayed at the same contrast levels. Top is FITC stained B220 B cell follicle region. Bottom is unlabelled (not B cell follicle) region. (b) Autofluorescence signal of B cell follicle region imaged in red in a control with no AF647 labelled chemokine and segmentation of this image. Scale bars 1μm.

## Supplementary video legends

**Supplementary Video 1: BSA-AF647 in PBS buffer. Whilst some particles are tracked, many diffuse further than the tracking radius in each frame, and cannot be tracked. Video after 260ms of imaging, slowed 10x. Tracked molecules are overlaid with a white line, scale bar 1 μm.**

**Supplementary Video 2: BSA-AF647 in 10% Ficoll 400, after 350ms of imaging, slowed 10x.The increase in viscosity by a factor of 5.6 compared to buffer alone allows the particles to be tracked. Tracked molecules are overlaid with a white line, scale bar 1 μm.**

**Supplementary Video 3: CCL19-AF647 diffusion in collagen, showing immobile population and stepwise photobleaching, slowed 10x. Tracked molecules are overlaid with a white line, scale bar 1 μm.**

**Supplementary Video 4: CXCL13-AF647 diffusion in collagen, showing members of both the mobile and immobile populations, slowed 10x. Tracked molecules are overlaid with a white line, scale bar 1 μm.**

**Supplementary Video 5: CXCL13-AF647 in lymph node tissue section. Tracks corresponding to both extracellular matrix and mobile chemokine are seen. 100ms of imaging is shown, slowed 10x. Tracked molecules are overlaid with a white line, scale bar 1 μm.**

**Supplementary Video 6: Heparan sulfate immobilised CCL19-AF647, showing a predominantly immobile population undergoing photoblinking behaviour, slowed 10x. The first frames after laser illumination are included. Tracked molecules are overlaid with a white line, scale bar 1 μm.**

**Supplementary Video 7: Heparan sulfate immobilised CXCL13-AF647, showing a predominantly immobile population, slowed 10x. Tracked molecules are overlaid with a white line, scale bar 1 μm.**

